# Establishment of Transgenic Fluorescent Mice for Screening Synaptogenic Adhesion Molecules

**DOI:** 10.1101/2022.07.20.500799

**Authors:** Lei Yang, Jingtao Zhang, Yanning Zhang, Li Wang, Xiaotong Wang, Shanshan Wang, Ke Li, Mengping Wei, Chen Zhang

**Affiliations:** School of Basic Medical Sciences, Beijing Key Laboratory of Neural Regeneration and Repair, Advanced Innovation Center for Human Brain Protection, Capital Medical University, Beijing 100069, China; Chinese Institute for Brain Research, Beijing 102206, China; Peking-Tsinghua Center for Life Sciences, Academy for Advanced Interdisciplinary Studies, Peking University, Beijing 100871, China

**Keywords:** synapse formation, Syt1-TDT transgenic mice, artificial synapse formation assay

## Abstract

Synapse is the fundamental structure for neurons to transmit information between cells. The proper synapse formation is crucial for developing neural circuits and cognitive functions of the brain. The aberrant synapse formation has been proved to cause many neurological disorders, including autism spectrum disorders and intellectual disability. Synaptic cell adhesion molecules (CAMs) are thought to play a major role in achieving mechanistic cell-cell recognition and initiating synapse formation via trans-synaptic interactions. Although several synaptic CAMs, such as neurexins, neuroligins, SynCAMs, and LRRTMs, have been identified as synaptogenic molecules, these molecules so far as we know cannot fully explain the mechanism of synapse formation. There should be other synaptogenic adhesion molecules that remain undiscovered. Artificial synapse formation (ASF) assays, the commonly used method for screening synaptogenesis molecules, is time-consuming and labor-intensive due to the long-lasting immunostaining step. To skip this step, we generated synaptotagmin 1-tdTomato (Syt1-TDT) transgenic mice by inserting the tdTomato-fused synaptotagmin 1 coding sequence into the genome of C57BL/6J mice. In the brain of Syt1-TDT transgenic mice, the Syt1-TDT signals were widely observed in different areas. In the cultured hippocampal neurons, the Syt1-TDT signals merged with several synaptic markers, suggesting the well synaptic localization of Syt1-TDT. Compared to the wild-type (WT) mouse neurons, cultured hippocampal neurons from Syt1-TDT transgenic mice presented normal synaptic neurotransmission. In ASF assays, neurons from Syt1-TDT transgenic mice could form synaptic connections with HEK293T cells expressing neuroligin2, LRRTM2, and Slitrk2 without immunostaining. Therefore, our work suggested that the Syt1-TDT transgenic mice with the ability to label synapses by tdTomato will be a convenient tool for screening synaptogenic molecules.

## Introduction

Synapses, consisting of the presynaptic membrane, the synaptic cleft, and the postsynaptic membrane, are the intercellular connections that rapidly transmit information from presynaptic neurons to postsynaptic cells in a point-to-point manner [1]. Synapse formation is an essential stage in brain development that begins during embryonic/postnatal development, and lasts throughout life [2]. The proper formation of synapses is crucial for neural circuits’ formation and cognitive function development, and abnormal synapse formation can cause many neurological disorders, including autism spectrum disorders (ASD) and mental retardation [3-6].

Synaptic cell adhesion molecules (CAMs) are molecules that act as a key role in initiating synapse formation through trans-synaptic interactions, and were originally proposed to enable mechanistic cell-cell recognition [7-9]. A variety of CAMs have been shown to initiate synapse formation, anchor and organize the precise alignment of the pre- and postsynaptic sides of a synapse, and enable enhanced short- and long-term synaptic plasticity of synaptic transmission [1]. For example, in vitro experimental studies have shown that neuroligins (NLGs) can trigger the re-formation of presynaptic structures. Non-neuronal cells overexpressing NLGs can induce presynaptic morphological and functional differentiation upon contact with axons [10]. The overexpression of neuroligin 1 (NLG 1) and neuroligin 2 (NLG 2) in vitro leads to the aggregation of presynaptic vesicles at both glutamatergic and GABAergic synapses [11]. Leucine-rich-repeat transmembrane neuronal proteins (LRRTMs) are a family of transmembrane proteins that induce presynaptic differentiation of contact axons, and all four LRRTM family members have synaptogenic activity [12]. The knockdown of LRRTM2 expression is associated with a decrease in the number of excitatory synapses [13], while overexpression leads to an increase [14]. Overexpression of netrin-G Ligands (NGLs), which are detected mainly at the postsynaptic sites of excitatory synapses and interact with postsynaptic density 95 (PSD95), facilitates pre- and postsynaptic differentiation, while the knockdown of NGLs leads to a decline in the number and function of excitatory synapses. Overexpression of NGL-2 in rat neurons increases the number of PSD95-positive dendrites, and direct accumulation of NGL-2 on the dendritic membrane surface triggers excitatory postsynaptic protein aggregation [15, 16], thereby demonstrating the impact of NGL-mediated adhesion effects in synapse formation.

However, very few CAMs have been identified compared to the large number of existing synapses. Knocking out these CAMs does not completely prevent neuronal synapses formation. For instance, in vivo studies on NLG1-NLG2-NLG3 triple knockout mice showed that the knockout of NLG1-NLG2-NLG3 did not alter the number of synapses in the brain, even though the mice were died due to the reduced GABAergic/glycinergic and glutamatergic synaptic transmission [17]. The number of pre-synaptic marker bassoon in hippocampus showed normal in LRRTM1-deletion mice [18]. No alteration in the density of PSD95 was detected in the netrin-G1 and netrin-G2 knockout mice’s hippocampus [19]. Therefore, there should be molecules that contribute to synapse formation remained undiscovered.

Up to now, a large-scale screen for synaptogenic CAMs remains difficult to implement. A study identifies gene transcripts encoding proteins expressed by postsynaptic neurons that potentially initiate the synaptic contacts’ formation by performing a genome-wide, expression analysis of chick ciliary ganglion at the different stages of synapse formation [20]. This screening method is comparatively complex and does not allow for direct observation of synapse formation. Another method for screening CAMs is the artificial synapse formation (ASF) assay [10]. It works by co-culturing non-neuronal cells, such as HEK293T cells, which is overexpressed by certain genes, with neurons. It can be observed that the generated synapses wrap around non-neuronal cells when the gene overexpressed has the synaptogenesis function. Many synaptogenic CAMs have been found to accumulate synapse around the non-neuronal cells in ASF experiments, including neurexins, neuroligins, SynCAMs, LRRTMs, latrophilins, protein tyrosine phosphatase receptor type O (PTPRO), and others [10-12, 21-24]. However, ASF assay still requires a lot of time for immunohistochemically staining, making it difficult to perform large-scale screening.

Synaptotagmin 1 (Syt1), a fast calcium sensor on synaptic vesicles [25, 26], begins to be strongly expressed on embryonic day 18 and continues to increase until reaching stable levels in adulthood (postnatal day 60) in many brain regions [27]. Syt1 is the most abundant synaptotagmin isomer in synaptic vesicles [28]. In the hippocampus, Syt1 is abundant in the stratum oriens, stratum radiatum, and stratum lacunosum moleculare [29]. In this study, we fused tdTomato (TDT) to the end of the Syt1 protein, and used PiggyBac transposon vector system to insert the synaptotagmin1-tdTomato (Syt1-TDT) gene into C57BL/6J mice allete. The tdTomato signals were widely observed in the brain of Syt1-TDT transgenic mice, and colocalized with synaptic markers. The transgenic mice can also be used for ASF experiments in order to visualize synapse by fluorescence microscopy without immunostaining. Therefore, the Syt1-TDT transgenic mice will be a valuable tool for large scale screening of CAMs.

## Results

### Syt1-TDT transgenic mouse expresses wildly distributed fluorescent signals in synapse-rich regions of the brain

We used fertilized eggs microinjection with PiggyBac vectors carrying Syt1-TDT to generate transgenic mouse line (C57BL/6J background) with insertions of tdTomato-fused synaptotagmin 1 into a mouse allele (Figures 1A). In the PiggyBac vector, there were two PiggyBac ITRs on either side of Human ubiquitin C (UBC) promoter-Kozak-Syt1-TDT-polyA in order to facilitate transposon-mediated transgene integration. To verify the successful construction of a Syt1-TDT expression vector, we digested the targeting vectors with restriction enzymes (Figures 1A and 1B). According to the schematic diagram of the linearized expression vector, the fragment sizes we obtained after enzyme digestion of the expression vectors should be AvaI: 4.7/1.7/0.7/0.5 kb (Figure 1B, left); AflII/NcoI: 4.0/1.3/1.1/0.7/0.4/0.2 kb (Figure 1B, middle); NotI: 7.6 kb (Figure 1B, right), which was consistent with our design. After a successful construction, the Syt1-TDT expression vector was co-injected with transposes into fertilized eggs from C57BL/6J mice. The offspring that carry the desired PiggyBac transgene were identified and selected by polymerase chain reaction (PCR).

**Figure 1.**
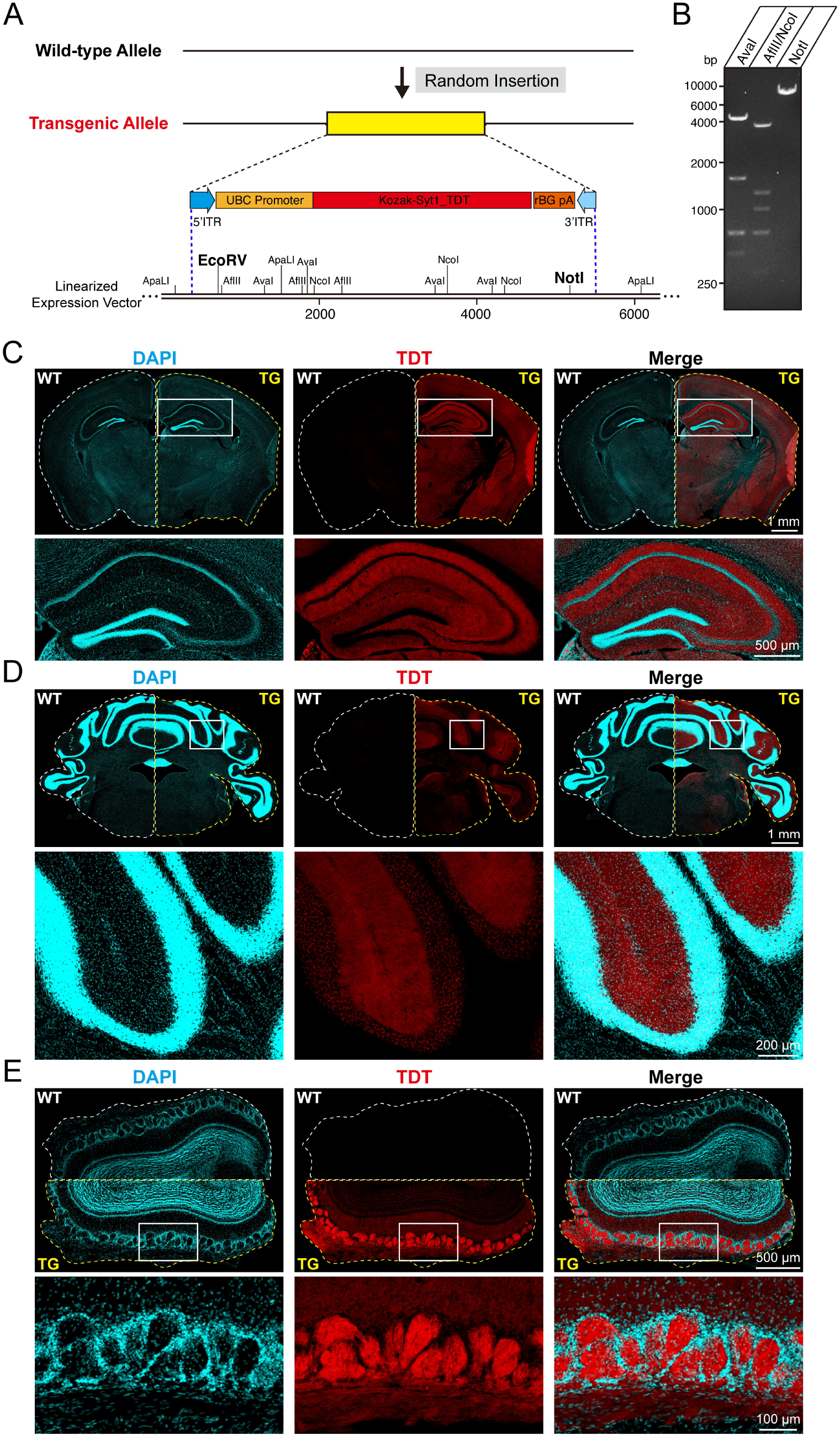
Construction of Syt1-TDT expression vector and transgenic fluorescent mice. (A) Schematic diagram of expression vector for generating Syt1-TDT transgenic mice. (B) The expression vector was digested by restriction enzymes for confirmation purposes. The size of enzyme digestion band (kb): (left) AvaI: 4.7/1.7/0.7/0.5; (middle) AflII/NcoI: 4.0/1.3/1.1/0.7/0.4/0.2; (right) NotI: 7.6. (C-E) Representative fluorescent images in different brain areas obtained from the Syt1-TDT transgenic mice and WT mice (8 weeks). (C) Hippocampus and cortex; (D) cerebellum; (E) olfactory bulb. All nuclei were stained with DAPI (cyan). **Figure 1-Source data** : the original gel of panel B.

To identify the distribution of Syt1-TDT signals in the brain of transgenic mice, we made frozen brain slices from 8-week-old Syt1-TDT transgenic mice and WT mice, and the brain slices were stained with DAPI to visualize the cell nucleus. We observed that Syt1-TDT was widely expressed in the hippocampus, cortex, cerebellum, and olfactory bulb in transgenic mice, but not in WT mice (Figures 1C, 1D, and 1E). In the hippocampus of transgenic mice, the signals of Syt1-TDT were relatively bright in the stratum oriens and stratum radiatum of CA1-CA3 and stratum lucidum of CA3, while the signals were absent in the stratum pyramidale (Figure 1C). In the cerebellum, the signals were stronger in the granular layer compared with molecular layer (Figure 1D). In the olfactory bulb, the Syt1-TDT showed bright signals in the glomerular layer (Figure 1E). Thus, our results suggested a wildly distribution of Syt1-TDT in different brain area, especially synapse-rich regions.

### Syt1-TDT signals colocalize with synaptic markers in cultured neurons

To observe whether the signals of Syt1-TDT localized well to the synaptic site, we performed immunostaining of synaptic markers in cultured hippocampal neurons from Syt1-TDT transgenic mice. We could observe strong tdTomato fluorescence in the cultured neurons which co-localized with the staining of synapsin (Figure 2A). Furthermore, we performed the immunostaining of pre- and post-synaptic markers, including vesicular glutamate transporter 1 (vGlut1), glutamic acid decarboxylase 65 (GAD65), PSD95, Homer1, and Gephyrin. Our results showed that the Syt1-TDT signals co-localized well with these synaptic markers (Figure 2B). The co-localization rates were identified as percentage of signal A colocalized with signal B in total signal A (A-B): Synapsin-TDT: 70.55 ± 1.447%; TDT-Synapsin: 48.55 ± 1.851%; vGlut1-TDT: 61.09 ± 1.517%; TDT-VGlut1: 52.88 ± 1.456%; GAD65-TDT: 52.08 ± 2.442%; TDT-GAD65: 44.43 ± 2.096%; PSD95-TDT: 68.50 ± 2.397%; TDT-PSD95: 62.38 ± 2.433%; Homer1-TDT: 61.00 ± 2.127%; TDT-Homer1: 62.13 ± 1.703%; Gephyrin-TDT: 48.87 ± 2.466%; TDT-Gephyrin: 39.36 ± 1.665% (Figure 2C). These results indicate that Syt1-TDT signals localize well to the synaptic site in the neurons from transgenic mice.

**Figure 2.**
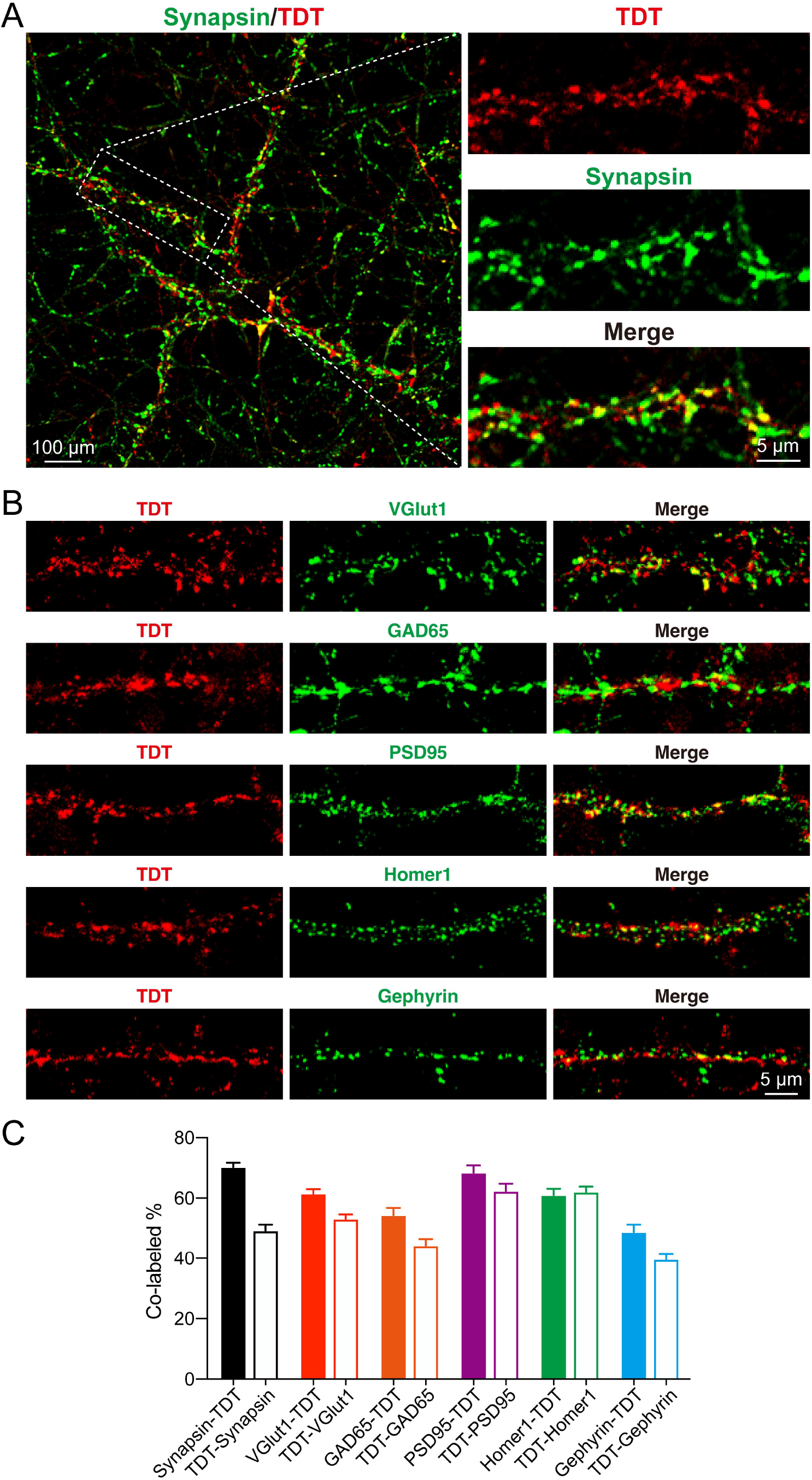
Syt1-TDT signals localize to the synaptic site in cultured neurons. (A-B) Representative images of hippocampal neurons from Syt1-TDT transgenic mice stained with antibodies against synapsin, Vglut1, GAD65, PSD95, Homer1 and Gephyrin at DIV14. (C) Summary graphs of co-labeling rate of Syt1-TDT with synapsin, Vglut1, GAD65, PSD95, Homer1 and Gephyrin (Synapsin: n=98/3, VGlut1: n=90/3, GAD65: n=71/3, PSD95: n=61/3, Homer1: n=81/3, Gephyrin: n=69/3). All summary graphs show the mean ± standard error of the mean (SEM). **Figure 2-Source data**: Numerical data corresponding to the graph in panel C

### Neurons from Syt1-TDT transgenic mice show normal excitability and synaptic transmission

To further explore whether the insertion of Syt1-TDT affects the electrophysiological properties and synaptic neurotransmission of neurons in transgenic mice, we performed patch clamp recordings on the cultured hippocampal neurons from Syt1-TDT transgenic mice and WT mice. We recorded the action potentials in these neurons and found that the amplitude, frequency, and half width of spontaneous action potentials in the neurons of Syt1-TDT transgenic mice are not significantly different with the WT mice (Amplitude: WT: 84.32 ± 2.930 pA, TG: 78.99 ± 3.325 pA; Frequency: WT: 0.4843 ± 0.1379 Hz, TG: 0.5424 ± 0.1391 Hz; Half width: WT: 2.178 ± 0.2616 ms, TG: 2.301 ± 0.2123 ms) (Figure 3C). These results demonstrate that insertion Syt1-TDT does not affect the excitability of neurons in transgenic mice.

**Figure 3.**
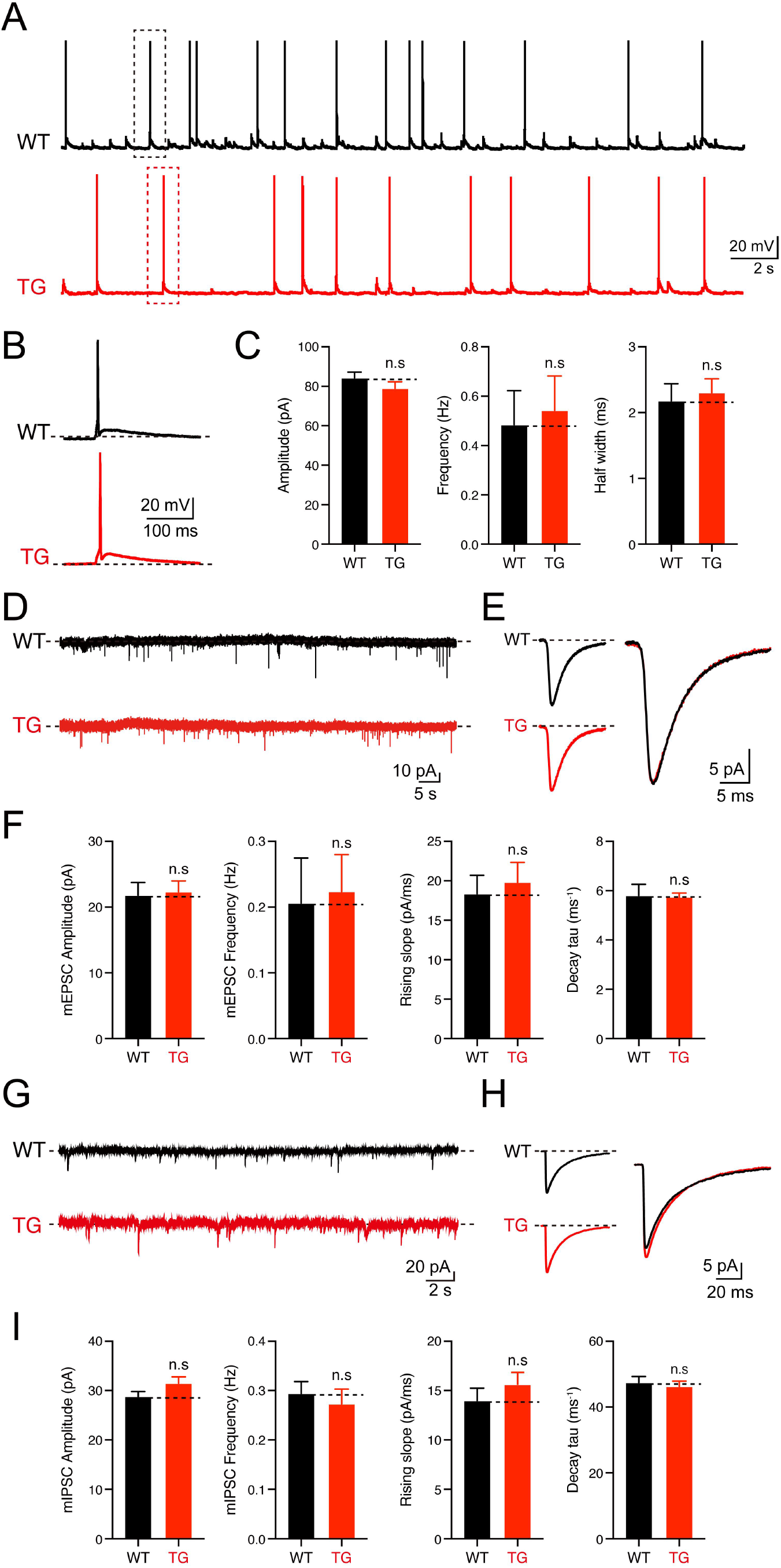
No significant changes of electrophysiological properties were observed in neurons of Syt1-TDT transgenic mice. (A-C) The spontaneous action potentials were recorded in hippocampal neurons of WT mice and Syt1-TDT transgenic mice (TG) at DIV 13-15. (A) representative traces of action potential; (B) representative traces of single action potential; (C) summary graphs of the amplitude (Left), frequency (Center) and half widths (Right). (WT: n = 43/3; TG: n = 45/3, amplitude: p=0.235, frequency: p=0.768, half widths: p=0.716). (D-F) The mEPSCs were recorded in hippocampal neurons of WT mice and Syt1-TDT transgenic mice in 1 μM tetrodotoxin (TTX) and 0.1 mM picrotoxin (PTX) at DIV 13-15. (D) representative traces of mEPSCs; (E) representative traces of normalized traces of mEPSCs; (F) summary graphs of the frequency (Left), amplitude (Center Left) of mEPSCs and the rising slope (Center Right), decay τ (Right) of normalized traces of mEPSCs. (WT: n = 21/3; TG: n = 26/3, amplitude: p=0.837, frequency: p=0.842, rising slope: p=0.681, decay τ: p=0.894). (G-I) The mIPSCs recordings in hippocampal neurons of WT mice and Syt1-TDT transgenic mice in 1 μM tetrodotoxin (TTX) and 10 μM CNQX at DIV 13-15. (G) representative traces of mIPSCs; (H) representative traces of normalized traces of mIPSCs; (I) summary graphs of the frequency (Left), amplitude (Center Left) of mIPSCs and the rising slope (Center Right), decay τ (Right) of normalized traces of mIPSCs. (WT: n = 59/4; TG: n = 59/4, amplitude: p=0.102, frequency: p=0.578, rising slope: p=0.355, decay τ: p=0.654). All summary graphs show the mean ± standard error of the mean (SEM); statistical comparisons were made by two-tailed unpaired t-test (n.s, not significant). **Figure 3-Source data**: Numerical data corresponding to the graph in panel C, F, I

We also examined the excitatory and inhibitory synaptic transmission mediated by AMPA receptors (AMPARs) or GABA receptors (GABARs) by recording the miniature excitatory postsynaptic currents (mEPSCs) and miniature inhibitory postsynaptic currents (mIPSCs). Our results shows that both the amplitude and the frequency of mEPSCs and mIPSCs of the Syt1-TDT transgenic mice neurons are similar to the WT mice (mEPSCs: Amplitude: WT: 21.97 ± 1.949 pA, TG: 22.50 ± 1.670 pA; Frequency: WT: 0.2078 ± 0.06898 Hz, TG: 0.2255 ± 0.05640 Hz; mIPSCs: Amplitude: WT: 28.83 ± 7.330 pA, TG: 31.48 ± 9.982 pA; Frequency: WT: 0.2942 ± 0.02385 Hz, TG: 0.2728 ± 0.03020 Hz) (Figures 3F and 3I). An analysis of the rising slope and decay time course of the mEPSCs and mIPSCs (Figures 3F and 3I) shows no significant differences between Syt1-TDT transgenic mice and WT mice, indicating that the expression of Syt1-TDT did not influence the kinetics of mEPSCs and mIPSCs. Thus, our results demonstrate that insertion of Syt1-TDT does not affect basal synaptic transmission.

### Neurons from Syt1-TDT transgenic mice can be used for screening CAMs in an artificial synapse formation assay

To test whether Syt1-TDT transgenic mice could be used for screening CAMs in an artificial synapse formation assay, we performed a co-culture assay between neurons from Syt1-TDT transgenetic mice and HEK293T cells transfected with NLG 2 and green fluorescent protein (GFP). Neuroligin 2 promotes synapse formation in ASF assays which can be used as a positive control [10, 24]. After being co-cultured for 36-48 hours, the cells were stained and observed using confocal microscopy. We observed a strong accumulation of tdTomato signals around the transfected HEK293T cells (indicated by GFP). Additionally, the accumulated signals overlapped highly with the staining signals of synapsin proteins (Figure 4A).

**Figure 4.**
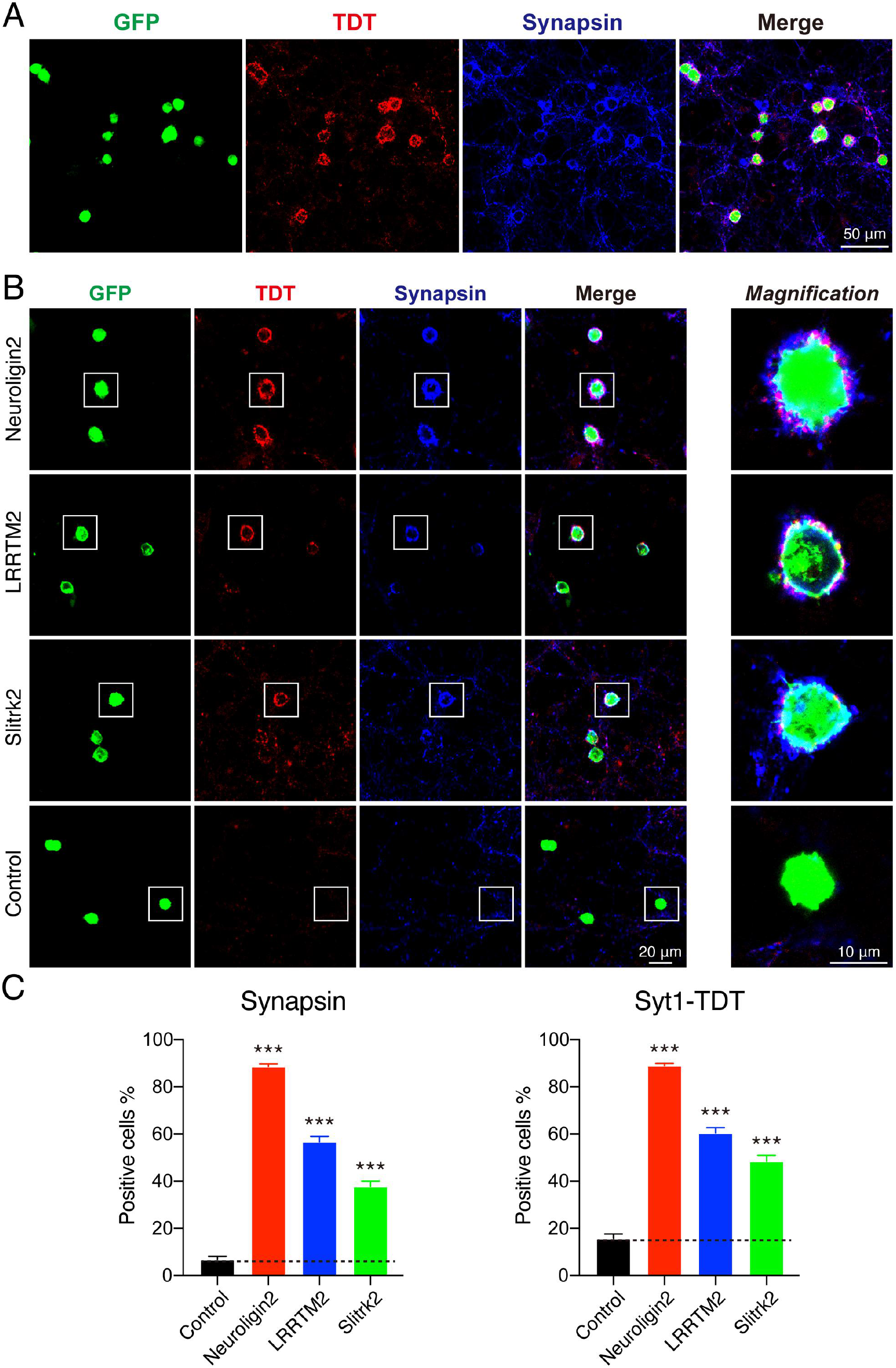
Syt1-TDT transgenic mouse hippocampal neurons can be used in artificial synapse formation experiments to screen for synaptogenic molecules. (A) Representative fluorescent images of co-cultured neurons from Syt1-TDT transgenic mice with HEK293T cells transfected with plasmid encoding neuroligin 2 and GFP, which are also stained with synapsin antibody. (B) Representative fluorescent images of co-cultured neurons from Syt1-TDT transgenic mouse with HEK293T cells transfected with plasmid encoding neuroligin 2, LRRTM 2, Slitrk 2 and GFP. (C) Summary graphs of the percentage of synapse-positive HEK293T cells from the panel B. (Synapsin: Control: n = 54 cells out of 854 cells/3 cultures; Neuroligin 2: n = 1417 cells out of 1599 cells/3 cultures; LRRTM 2: n = 692 cells out of 1259 cells/3 cultures; Slitrk 2: n = 374 cells out of 1032 cells/3 cultures. TDT: Control: n = 148 cells out of 854 cells/3 cultures; Neuroligin 2: n = 1411 cells out of 1599 cells/3 cultures; LRRTM2: n = 740 cells out of 1259 cells/3 cultures; Slitrk 2: n = 486 cells out of 1032 cells/3 cultures). All summary graphs show the mean ± standard error of the mean (SEM); statistical analysis was made by one-way ANOVA (***p < 0.001). **Figure 4-Source data**: Numerical data corresponding to the graph in panel C

To further confirm that neurons from Syt1-TDT transgenic mouse can be used in ASF assays to screen synaptogenic molecules, we also tested other synaptogenic CAMs in ASF assays using Syt1-TDT neurons. Our results showed that NLG2, LRRTM2, and Slitrk2 enabled the formation of synaptic connections between transfected HEK293T cells and neurons from Syt1-TDT transgenic mice. We compared the percentage of synapse-positive HEK293T cells calculated by either tdTomato’s signals or the stained synapsin signals as previously report [22], and found that both of signals had the ability to labeling the synapse accumulated around the HEK293T cells (synapsin: Control: 6.802 ± 1.329 %, NLG 2: 88.65 ± 1.077 %, LRRTM2: 56.69 ± 2.234 %, Slitrk2: 37.82 ± 2.216 %; TDT: Control: 15.70 ± 1.949 %, NLG 2: 88.94 ± 0.9419 %, LRRTM2: 60.48 ± 2.186 %, Slitrk2: 48.47 ± 2.469 %) (Figures 4B, 4C). According to the above results, we thought that neurons Syt1-TDT transgenic mice are suitable for screening CAMs in an ASF assay.

## Discussion

In our study, we generated transgenic mice that label synapses by expressing fluorescent proteins to skip the immunofluorescence staining step during synaptogenic adhesion molecules screening. We chose Syt1 as a media to transport tdTomato to the synapse by fusing tdTomato to the C-tail of Syt1. We identify the synaptic localization and no effect on synaptic transmission of Syt1-TDT. We demonstrated its availability for ASF assay, which is useful for synaptogenic adhesion molecule screening.

We designed the Syt1-TDT expression vector and injected it into the fertilized eggs of C57BL/6J mice in order to insert the Syt1-TDT gene into the genome of C57BL/6J mice to create transgenic fluorescent mice. Due to random insertion, we do not know the actual insertion site and its number. However, based on the microscopic imaging of frozen sections of Syt1-TDT transgenic mice brain (Figures 1C, 1D, and 1E), Syt1-TDT signals were observed in the hippocampus, cortex, cerebellum, and olfactory bulb, and the areas of strong signals were in keeping with previous studies on the distribution of Syt1 in the brain [29, 30]. The co-labeling of synaptic markers in the cultured hippocampal neurons with Syt1-TDT signals (Figures 2A, 2B, and 2C) further confirmed that fluorescence of Syt1-TDT is well localized to synapses. Although we usually thought that Syt1 is located at presynaptic site as a fast calcium sensor on synaptic vesicles, we found that Syt1-TDT signals also have a high co-labeling rate with postsynaptic protein markers (Figure 2C). We proposed that may be due to:1. We cannot distinguish the pre- and post-synaptic site because of the limitation of resolution of confocal microscope; 2. The presence of syt1 at the post-synaptic site. A recent study has shown that Syt1 is also expressed in hippocampal postsynaptic spines [31]. Even so, our results suggested that Syt1-TDT is good for labeling synapse.

Overexpression of synaptic proteins were reported to affect the synaptic function. For instance, overexpression of synaptic vesicle protein synapsin caused the decrease of basal transmission and increase of homosynaptic depression [32]. Overexpression of post-synaptic scaffolding protein SAP97 caused the modifications of spine and synapse morphology [33]. Overexpression of synaptic CAM neuroligins caused the change of excitatory and inhibitory synapse properties [34]. We found that the electrophysiological properties of Syt1-TDT transgenic mice neurons were not significantly different from WT mice neurons (Figures 3A-3I). This provides us with better conditions for synaptogenic adhesion molecules screening with fewer effects on synaptic transmission. This also offers the possibility for Syt1-TDT transgenic mice to be used for labeling synapse to facilitate the study of electrophysiological recording and imaging in vivo.

In a previous work Synaptophysin-YFP was used to labeling neurites in pontine or vestibular explants-neuronal granule cells co-culture assay [35], it provided us an idea that the synaptic labeling neurons could be used for visualize artificial synapse formation between neurons and non-neuronal cells. Syt1-TDT transgenic mice neurons were able to label positive HEK293T cells expressing synaptogenic molecules (NLG2, LRRTM2, and Slitrk2) without immunostaining (Figures 4A, 4B, and 4C). Because NLG2, LRRTM2, and Slitrk2 are mainly expressed at the postsynaptic site, they can be recognized by presynaptic terminals marked by Syt1-TDT to form intact synapses. Even though it has been suggested that Syt1 is also expressed postsynaptic site, it remains unclear to us whether Syt1-TDT transgenic mice neurons can be used for screening synaptogenic molecules located at presynaptic site. In order to establish a complete screening system, we have also started to construct the transgenic fluorescent mice model that expresses fluorescent proteins and specifically transport them to postsynaptic spines.

The ASF assays using Syt1-TDT transgenic mice to screen for synaptogenic molecules have skipped the necessity of immunostaining. However, we still can only test the synapse accumulation around HEK293T cells which are transfected with the same plasmid. It requires large numbers of neurons and HEK293T cells for large-scale screening. To optimize the screening process for large-scale and high-throughput screening, we are examining the co-culture of neurons with multiple HEK293T cells transfected with different plasmids in the same well. The synapse-accumulated HEK293T cells in the co-culture system can be separated by cell digestion, single-cell picking, and single cell sequencing can be used to identify the genotype the picked cells. If it works, it will largely improve the throughput of screening in ASF assay.

## Materials and Methods

### Construction of Syt1-TDT expression vector and generation of Syt1-TDT transgenic mouse

The PiggyBac vector and UBC promoter (human ubiquitin C promoter) were chosen to construct the Syt1-TDT expression vector. In the PiggyBac vector, there are two PiggyBac ITRs located at the sides of the “UBC promoter-Kozak-Syt1-TDT-polyA” cassette in order to promote transposes mediated transgene integration. The constructed vectors were co-injected with transposes into fertilized eggs from C57BL/6J mice. The offspring were identified by PCR to select those carrying the required PiggyBac transgene. Counter screening was performed on positive founder mice for transposes. The Cyagen Biosciences (Guangzhou) Inc. conducted all of the processes.

### Animals

This study used C57BL/6J WT mice and Syt1-TDT transgenic mice (P0-P56). Animals were housed at room temperature (RT) 20 ± 2°C, with a 12-hour light-dark cycle, air circulating, and unrestricted access to food and water. All procedures and experiments were performed in conformance with the ethical regulations that the Animal Experimentation Committee of Capital Medical University established.

### Genotyping

Genomic DNA was isolated from the tail piece. This was done by taking an appropriate amount of rat tail (about 0.5 cm long) and placing it in a 1.5 ml EP tube. We then added 200 μl 50 mM NaOH and heat in a 100°C water bath for 30 min for lysis. After vortex mixing, 16 μl 1 M tris-HCL was added to end lysis. For PCR analysis, primers were designed according to the Syt1-TDT expression vector (primer F: TAGGGTAGGCTCTCCTGAATCGAC; primer R: ATTCATAAACTTCTGCTTCAGCTTG). The PCR was performed with the 2 × Taq Plus Master Mix II (Vazyme, P213-01) and according to the manufacturer’s instructions. The PCR products carried out electrophoresis on 1.5% agar gel. The amplification product was 396 bp.

### Primary hippocampal neuronal culture

Neurons for experiments were obtained from postnatal day 0 C57BL/6J mice’s hippocampus. The mice’s brains were isolated, and the hippocampus were dissected on ice. After washing with DMED (Gibco, 11995-065), the hippocampus was placed in 0.25% trypsin (Sigma, T4049) and digested at 37°C for 12 min. We added DMEM to end the digestion, and to count after blowing and mixing. Neurons were plated on glass coverslips (diameter 12 mm) that were coated with poly-D-lysine (Gibco, A38990401) at a density of 80,000 neurons per coverslip and cultured in Neurobasal Plus Medium (Gibco, A35829-01) containing 2% B-27 Plus Supplement (Gibco, A3592801) at 37°C in 5% CO2 in a thermostatic cell incubator.

### Immunofluorescence staining of neurons

The cultures were fixed with 4% paraformaldehyde in phosphate-buffered saline (PBS, pH 7.4) for 10 min. After washing with PBS, the cultures were blocked and permeabilized with Blocking Buffer (0.3% Tween 20, 5% skimmed milk, and 2% goat serum in PBS). Cultures were incubated overnight with primary antibodies [anti-synapsin (1:20,000; homemade), anti-vGLUT1 (1:2,000; Synaptic Systems, 135 302), anti-GAD65 (1:5,000; Sigma, G5638), anti-PSD95 (1:2000; NeuroMab, 75-028), anti-Homer1 (1:1000; Synaptic Systems, 160 011), or anti-Gephyrin (1:1000; Synaptic Systems, 147 011)] diluted in Blocking Buffer. After washing three times for 5 min in PBS, cultures were incubated for 30 min at room temperature with secondary antibodies [Goat anti-Rabbit IgG (H+L) Highly Cross-Adsorbed Secondary Antibody, Alexa Fluor Plus 647 (1:200; Invitrogen, A32733) or Dylight 649, Goat Anti-Mouse IgG (1:200; Abbkine, A23610)]. Samples were placed in mounting medium (Southernbiotech, 0100-01) after washing them three times for 5 min in PBS and once in double distilled H2O (ddH2O).

### Immunofluorescence staining of mouse brain slices

Adult mice (transgenic and WT) were deeply anesthetized with tribromoethanol and then transcardially perfused with 4% paraformaldehyde in phosphate-buffered saline (PBS, pH 7.4). The brains were removed and fixed in the 4% paraformaldehyde in PBS for 24 hours. After tissue dehydration with 15% and 30% sucrose solution in PBS, brains were embedded with freezing embedding medium (optimum cutting temperature compound). The 40μm thick sections were cut on a cryo-cut cryostat microtome. The sections were permeabilized with 0.3% Triton X-100 in PBS. After washing for 5 min in PBS, sections were incubated in PBS containing 5 μg/ml DAPI (Beyotime, C1002) for 20 min in order to stain cell nuclei, washed with PBS, and placed in mounting medium.

### Electrophysiological recordings

Neurons at DIV 13-15 were taken for the experiment. Current-clamp recordings of primary hippocampal neurons were carried out with a MultiClamp 700A amplifier (Molecular Devices). Series resistance was compensated to 60-70%, and recordings with series resistances of > 20 MΩ were rejected. For AP recordings, neurons were patched and held in the current-clamp whole-cell configuration, and maintained in an external solution of 150 mM NaCl, 4 mM KCl, 10 mM HEPES, 2 mM CaCl2, 1 mM MgCl2, 10 mM glucose (pH 7.40, Osm 315 mOsm/kg). Microelectrodes (World Precision Instruments) were filled with internal solution, which contains 145 mM KCl, 5 mM NaCl, 10 mM HEPES, 5 mM EGTA, 4 mM MgATP, 0.3 mM Na2GTP (pH 7.25, Osm 305 mOsm/kg). Whole-cell voltage clamp recordings of primary hippocampal neurons were carried out using a MultiClamp 700A amplifier (Molecular Devices). Coverslips, seeded with primary hippocampal neurons, were kept in an external solution of 150 mM NaCl, 4 mM KCl, 2 mM CaCl2, 1 mM MgCl2, 10 mM HEPES, and 10 mM Glucose (pH 7.40, Osm 315 mOsm/kg). For mEPSC/mIPSC recordings, tetrodotoxin (1 µM) was added to the external solution in order to block the evoked synaptic responses. Microelectrodes (World Precision Instruments) were filled with an internal solution of 110 mM Cs-Methanesulfonate, 20 mM TEA-Cl, 8 mM KCl, 10 mM EGTA, 10 mM HEPES, 3 mM MgATP, and 0.3 mM Na2GTP (pH 7.3, Osm 275-285 mOsm/kg). The data were analyzed with Clampfit 10.2 (pClamp) and Igor 4.0 (WaveMetrics).

### HEK 293T cell culture and transfection

HEK293T cells were obtained from the Kunming Institute of Zoology, Chinese Academy of Sciences. They were cultured in COS medium (DMEM containing 10% FBS, 50 U/ml penicillin, 50 μg/ml streptomycin, 44.52 mM NaHCO3, pH 7.2-7.4) at 37°C in 5% CO2 in a thermostatic cell incubator. For passaging, when cells grow to 80% cell density, discard the old medium, and add PBS to wash away metabolic waste and floating dead cells. Add 0.05% of Trypsin solution, digest in the incubator for 2 min, and add 2 times the volume of COS medium to terminate the digestion. Centrifuge the cell suspension and add COS medium to resuspend the cells to culture. The HEK293T cells to be transfected were cultured in 24-well plates. After the cell density grew to about 80%, transfection was performed. Add plasmid and polyethylenimine (PEI; 1 mg/ml in ddH2O) to 35 µl OPTI. The PEI plasmid ratio was 3:1. The mixture was incubated at room temperature for 30 min and then added dropwise to HEK293T cell cultures. For one well of a 24-well plate, 1.2 μg of each plasmid and 0.3 μg of GFP was transfected.

### Artificial synapse formation

After 24 hours of transfection, the HEK293T cells were digested by adding 0.05% trypsin, placed at 37 °C for 1 minute, and the digestion was terminated by adding COS medium. The cell suspension was centrifuged, resuspended in Neurobasal Plus Medium (containing B-27), counted, and then added dropwise into hippocampal neuronal culture at DIV 9 in vitro (40,000/ coverslip), gently shaken, and placed at 37°C in 5% CO2 in a thermostatic cell incubator. After 36-48 hours of incubation, immunocytochemical assays were performed.

### Confocal imaging and Image Analysis

For immunofluorescence staining of neurons and mouse brain slices, images were captured using a confocal microscope (Olympus FV1000). Image J was used to analyze the co-labeling rate, and Matlab was used to analyze positive cells in the artificial synapse formation assay.

### Statistical analysis

Statistical analyses were conducted using GraphPad Prism 8.0.1 software for t-test. Results were displayed as mean ± standard error (SEM). Differences in means were accepted as significant if the results of the two-tailed unpaired t-test and one-way ANOVA were p < 0.05.

## Acknowledgments

This work was supported by grants from National Key R&D Program of China [2017YFA0105201]; the National Science Foundation of China [81925011, 92149304, 31900698, 32170954, 32100763]; The Youth Beijing Scholars Program (015), Support Project of High-level Teachers in Beijing Municipal Universities [CIT&TCD20190334]; Beijing Advanced Innovation Center for Big Data-based Precision Medicine, Capital Medical University, Beijing, China [PXM2021_014226_000026].

## Declaration of interests

The authors declare no competing interests.

## Data and Materials availability statement

We provide the catalog number of all the commercial materials in the Methods section The Syt1-TDT transgenic mice and the home-made synapsin antibody are available. If someone need, please contact the corresponding author. All the source data are available in Figure 1-4 -source data or in: Yang, Lei et al. (2022), Source data-Establishment of Transgenic Fluorescent Mice for Screening Synaptogenic Adhesion Molecules, Dryad, Dataset, https://doi.org/10.5061/dryad.dncjsxm30 Link: https://datadryad.org/stash/share/sW0OWEw5ubKTVIWQ8_kZdE0U6aUnmWZlOvXBh9×92Lc

## Notes

### Competing Interest Statement

The authors have declared no competing interest.

